# How sequence populations persist inside a genome

**DOI:** 10.1101/2020.06.25.170514

**Authors:** Hye Jin Park, Chaitanya S. Gokhale, Frederic Bertels

**Author notes:** These authors contributed equally to this work.

## Abstract

Compared to their eukaryotic counterparts, bacterial genomes are small and contain extremely tightly packed genes. Therefore, discovering a large number of short repetitive sequences in the genomes of Pseudomonads and Enterobacteria is unexpected. These sequences can independently replicate in the host genome and form populations that persist for millions of years. Here we model the interactions of intragenomic sequence populations with the bacterial host. In a simple model, sequence populations either expand until they drive the host to extinction or the sequence population gets purged from the genome. Including horizontal gene transfer does not change the qualitative outcome of the model and leads to the extinction of the sequence population. However, a sequence population can be stably maintained, if each sequence provides a benefit that decreases with increasing sequence population size. But concurrently, the replication of the sequence population needs to be costly to the host. Surprisingly, in regimes where horizontal gene transfer plays a role, the benefit conferred by the sequence population does not have to exceed the damage it causes. Together, our analyses provide a plausible scenario for the persistence of sequence populations in bacterial genomes. More importantly, we hypothesize a limited biologically relevant parameter range, which can be tested in future experiments.

---

Repetitive sequences can be found in most genomes. They are particularly abundant in eukaryotes, where often only a small proportion of the genome encodes for host proteins [1]. In contrast, about 90% of a typical bacterial genome encodes for host proteins [2]. The extragenic space is mostly taken up by rRNA, tRNA, transcription and translation promoters, repressors and terminators [3]. Yet, repetitive sequences can be found in the extragenic space of many bacteria [4].

Repetitive sequences were first identified in *Escherichia coli* in the early 1980s [5]. Then, due to their sequence characteristics, they were called REP sequences, short for **<u>r</u>**epetitive **<u>e</u>**xtragenic **<u>p</u>**alindromic sequences [6]. However, it was unclear if REP sequences fulfil a functional role in the host bacterium and if so what kind of function this might be. Numerous studies found REP sequences to be involved in different biological processes, for example in transcription termination, RNA stabilisation, gyrase and integration host factor binding, as well as nucleoid folding [5, 7–11]. However, whether the identified functions are locally co-opted, or common to all REP sequences and therefore able to explaining the presence of REP sequences in the bacterial genome, is not clear.

To determine whether a function is incidental or whether it can explain the persistence and emergence of an entire sequence class requires the understanding of the evolution of REP sequences. A study in *Pseudomonas fluorescens* SBW25 showed that REP sequences are not evolutionary relevant units [12], but a part of a larger replicative unit, called REPIN (**<u>REP</u>** doublets forming hairp**<u>ins</u>**). REPINs consist of two inverted REP sequences separated by a short and highly diverse spacer region. This arrangement allows REPINs to form hairpins in single-stranded DNA or RNA. REP singlets also exist, but these are usually decaying remnants of full-length REPINs. Furthermore, multiple studies simultaneously showed that there is a single transposase responsible for REPIN replication called RAYT (**<u>R</u>**EP **<u>a</u>**ssociated t**<u>y</u>**rosine **<u>t</u>**ransposase) [12–14].

RAYT transposases are single-copy genes that have been vertically inherited for millions of years [15]. Therefore RAYTs are domesticated transposases. Despite the domestication of the RAYT transposase by the bacterium, RAYTs have not lost their association to REPINs and seem to still actively replicate REPINs albeit at very low rates [16]. Although, the exact function of the RAYT transposase is unknown it is easily perceivable that formerly parasitic genes are domesticated by the host. It is much less clear how a population of replicating sequences can be maintained in a bacterial genomes over long periods of time.

Hence, here we want to explore what evolutionary conditions allow a sequence population to persist inside a genome. We show that if a sequence population does not confer a benefit to the host bacterium, it will either drive the bacterial population extinct or be lost from the bacterial population. Even horizontal gene transfer cannot explain the stable persistence of an intragenomic sequence population. However, persistence is possible when each sequence provides a small benefit to the host bacterium that decreases as the sequence number per genome increases. Interestingly, for high horizontal gene transfer rates, sequence populations can persist even if the caused harm outweighs the fitness benefit provided to the host. Together, our analyses provide testable hypotheses to explain the persistence of intragenomic sequence populations in bacteria.

## I. MATERIALS AND METHODS

### A. Local REPIN amplification rate λ

REPINs are often found in two or more tandem repeat copies [12, 17]. Hence, it is possible for REPINs to locally amplify and delete themselves. To estimate the rate at which local amplification and deletion happens in the genome we consulted mutation accumulation data from *Escherichia coli* MG1655 [18]. In this experiment the authors started 50 parallel mutation accumulation lines from a single *E. coli* MG1655 wild type clone. These 50 lines were grown on minimal medium and serially transferred about 220 times through single cell bottlenecks. Between bottlenecks the cells grew for about 28 generations. The final bacterial clones experienced about 6000 cell divisions from the start to the end of the experiment [18, 19]. At the end of the experiment the authors observed 277 single base pair substitutions across the 50 individual mutation accumulation lines and based on this data estimated a per genome mutation rate of 277(*substitutions*) / (6160(*generations*) × 50(*lines*)) ≈ 0.9 × 10^−3^.

Using the same logic, and further data from [20] we can estimate the local amplification rates of REPINs. Across the experiment they only observed a single large indel that involved REPINs and hence is relevant for the estimation of local REPIN amplifications and deletions (λ). This event occurred in M2M-85 (SRA accession number: SRR2169198) at position 4295870..4296434 in the *E. coli* MG1655 ancestor (Genbank accession number: U00096.3) and deleted five REPIN copies in a tandem cluster of six REPINs. From these numbers we can estimate the magnitude of the amplification rate λ the same way Lee et al. have done for the substitution rate. To focus on the rate per REPIN we have to also divide by the REPIN population size of *E. coli* MG1655 (224) to obtain λ = 1/(6160 × 50 × 224) ≈ 1.45 × 10^−8^.

### B. REPIN duplication rate *δ*

To determine REPIN duplication rates, we first identified the most common 21 to 25 bp long sequences in ten different Enterobacterial strains. For each of these highly abundant sequences, we determined the corresponding REPIN populations by recursively determining all sequences that differ in exactly one position from any already identified sequence [see [16] for more details]. By using the mutation-selection (or quasispecies) model, we inferred REPIN duplication rates as described in [16]. This model considers four mutation classes. The first mutation class only contains a single sequence, the master sequence. The second and third mutation classes contain all sequences that differ from the master sequence in exactly one and two positions, respectively. The last mutation class contains all sequences that differ in three or more positions to the master sequence. By assuming that the frequency distribution of the four mutation classes is in a steady-state, the REPIN duplication rate can be estimated for a constant mutation rate. Using this procedure, we obtained five duplication rates for the master sequence per bacterial strain, one for each sequence length. For each strain, we report the highest master sequence duplication rate. All estimated duplication rates are summarised in Table I.

**TABLE I.** Estimated duplication rates and REPIN population sizes

| Strain | Seq. length | Duplication rate ($\delta$) | REPIN pop. size ($r$) |
| --- | --- | --- | --- |
| *Salmonella enterica* ATCC 9150 | 24 | $5.2 \times 10^{-9}$ | 98 |
| *Citrobacter koseri* ATCC BAA-895 | 24 | $3.8 \times 10^{-9}$ | 323 |
| *Enterobacteriaceae bacterium* FGI 57 | 21 | $9.3 \times 10^{-9}$ | 150 |
| *Klebsiella variicola* 342 | 23 | $5.4 \times 10^{-9}$ | 91 |
| *Escherichia albertii* 07-3866 | 24 | $7.6 \times 10^{-9}$ | 226 |
| *Escherichia coli* K-12 MG1655 | 23 | $1.2 \times 10^{-8}$ | 224 |
| *Escherichia coli* B REL606 | 24 | $9.7 \times 10^{-9}$ | 220 |
| *Escherichia coli* UMN026 | 21 | $1.4 \times 10^{-8}$ | 159 |
| *Escherichia coli* UTI89 | 24 | $7.7 \times 10^{-9}$ | 137 |
| *Escherichia coli* 536 | 24 | $8.7 \times 10^{-9}$ | 158 |

## II. MODEL

Our main objective is to explore the conditions that would allow REPINs to persist in their bacterial host genome for millions of years or billions of bacterial generations. We begin by describing the dynamics of the hosts—the bacteria. We assume that bacteria grow near exponentially when the population size is small, and growth saturates when the population size is close to carrying capacity (i.e. logistic growth).

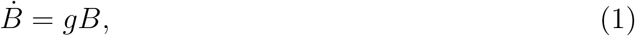

where *B* is defined as *B* = *n*/*K*, *K* is the population carrying capacity, and *n* is the number of bacteria in the population. *g* is defined as *g* =1 – *B*.

We can define bacterial subpopulations depending on the number of REPINs *r* each bacterium carries. The relative abundance of bacteria carrying *r* REPINs with respect to *K* is denoted by *b_r_* = *n_r_/K*. The bacterial pool is the sum of all bacteria with different numbers of REPINs, 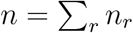. Hence *B* becomes

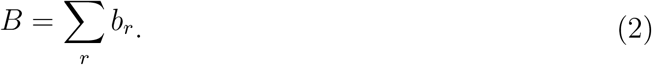

The number of bacteria carrying *r* REPINs changes not only because of the growth of the bacteria themselves but also due to the REPIN dynamics. For example, if a REPIN is deleted, the bacterium changes its state from *r* to *r* – 1, which happens with rate *w*_*r*, *r*–1_. Similarly if a REPIN successfully duplicates then we see the transition from *r* to *r* + 1, which happens at rate *w*_*r*, *r*+1_. The REPIN dynamics are sketched in Fig. 1. Altogether, the change in the relative bacterial abundance is captured by the following set of differential equations,

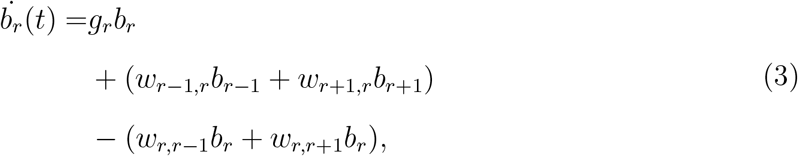

where *b_r_* = 0 for *r* < 0.

**FIG. 1.**
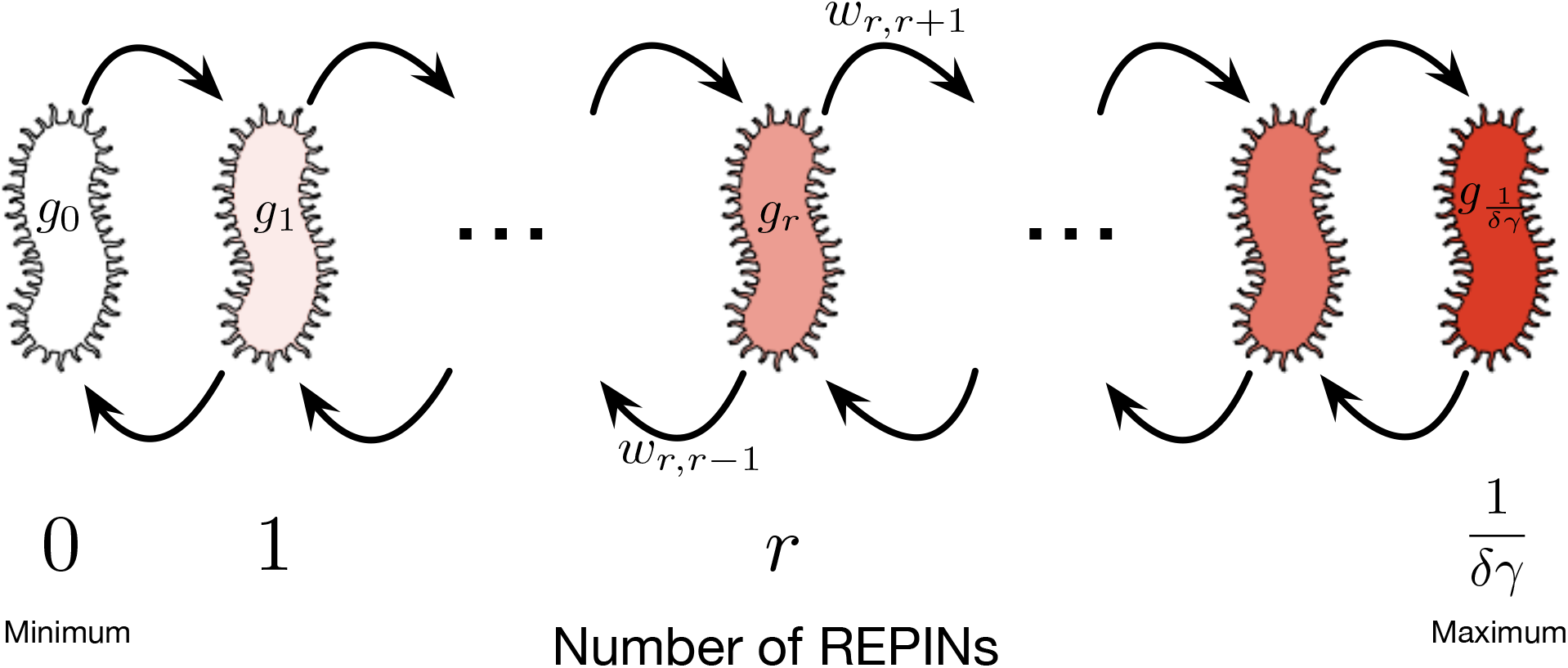
Modelling intragenomic sequence population in a bacterial population. Bacteria in the population only differ in the number of REPINs they contain. A bacterium with *r* REPINs gains a REPIN with rate *w*_*r*, *r*+1_ and loses a REPIN with rate *w*_*r*, *r*–1_. The gain and loss of REPINs depend on the parameter λ (random amplification and deletion of REPINs) and *δ* (REPIN duplication rate). The duplication rate *δ* also decreases the growth rate of each bacterium by *rδγ*, since with probability *γ* a bacterium will be killed after a duplication event. The minimum number of REPINs is zero, the upper REPIN population size limit for maintaining a viable bacterial population is given by *r* = 1/(*δγ*).

We connect growth and transition rates in the above equation with our observation in the previous section. The RAYT transposase duplicates REPINs by copying them into another location of the genome [12]. This duplication rate is denoted as *δ*. However, this type of duplication comes at a cost. Once a REPIN is copied into a gene, then the gene will be destroyed. If the gene is essential for bacterial survival, then the bacterium that carries the REPIN population, including the duplicated REPIN, will die. We denote *γ* as the fatality probability that a bacterium dies due to the duplication of a REPIN. Hence bacterial growth rate *g_r_* can be written as

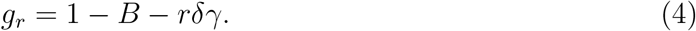

Additional to duplication by the RAYT transposase, our observation of REPINs in bacterial genomes suggests that REPINs may be able to reproduce locally. Local amplification and deletion of REPINs is probably mediated by the host replication machinery and not by the RAYT transposase [12, 21]. This mode of amplification and deletion is captured by including a birth rate *λ* and an equal death rate *λ* giving the transition rates

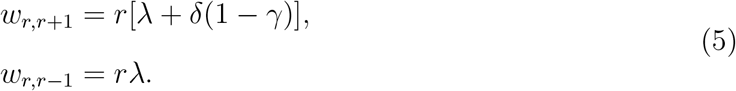

## III. RESULTS

### A. Simple replicating intragenomic sequence populations cannot persist in bacterial genomes

Our model describes a bacterial population in which each bacterium carries a certain number of REPINs *r*. If the number of REPINs does not affect the bacterial dynamics, meaning duplication does not induce the death of a bacterium, then the duplication process leads to an ever-increasing REPIN population.

However, every REPIN duplication can harm the bacterial growth rate. There is a chance *γ* that a REPIN duplication leads to the death of the bacterial host. This model will lead to two different outcomes depending on the parameter values and initial conditions. Either the REPIN population will go extinct in the bacterial population (*b*_0_ = 1, purple distribution in Fig. 2) or the REPIN population will grow uncontrolled and drive the bacterial population to extinction (*B* = 0, green distributions in Fig. 2).

**FIG. 2.**
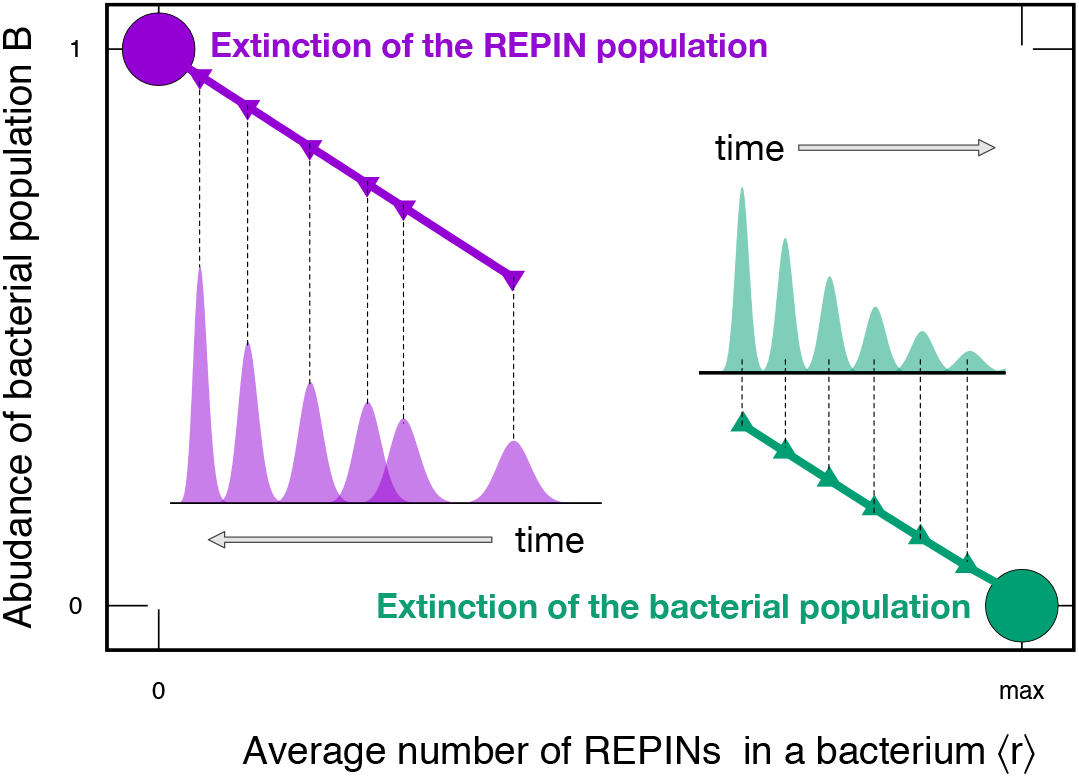
Base model showing the dynamics of the bacterial pool *B* and the average number of REPINs found in a bacterial genome 〈*r*〉. For different initial conditions, the dynamics show two different outcomes indicated by the purple and green circles. When the initial condition is close to the zero-REPIN state, the population follows the thick purple line leading to the extinction of the REPIN population inside the bacterial population. When starting with a large initial REPIN population size with small *γ*, the REPIN population size will increase. In contrast, the bacterial population size will decrease and eventually go extinct (green line). Each point marked with an arrow shows the distribution of the bacterial population *b_r_* at that time point. The figure shows simulation results for high parameter values: *δ* = 10^−3^, *λ* = 10^−5^ and *γ* = 0.2 for different initial distribution *b_r_*(*t* = 0). Results remain qualitatively the same for lower, more realistic parameter values.

For *γ* > 0 any duplication event can lead to the death of the bacterial host, and thus the fittest subpopulation is the population without REPINs. Bacteria devoid of REPINs have the highest growth rate. They cannot acquire REPINs in the absence of horizontal gene transfer. Hence, as soon as a fraction of bacteria loses all REPINs, REPINs will go extinct in the bacterial population. REPIN extinction usually happens when a population starts with small REPIN numbers or large *γ* values. When *γ* is large, bacteria where a REPIN duplicates are more likely to die than successfully increasing the REPIN number (purple distribution in Fig. 2).

Alternatively, the accumulation of REPINs can lead to the extinction of the bacterial population. The bacterial population will go extinct when large REPIN numbers accumulate, for example when *γ* is low (the bacterium is unlikely to die due to a duplication event). In this case, an increasing number of REPINs will lead to a decreasing number of bacteria. Thus eventually the entire population becomes extinct (green distribution Fig. 2).

We analytically prove that these two trivial scenarios are the only possible stable solutions of our model (Appendix B for detailed calculations). In either of the two scenarios the REPIN population will go extinct. Hence, our basic model does not explain what we observe in nature: an intragenomic sequence population that persists for millions of years.

### B. Horizontal gene transfer within a bacterial population cannot explain REPIN persistence

Horizontal gene transfer (*hgt*) has been shown to be essential to explain the persistence of selfish genetic elements [22, 23]. Although for REPINs there is no evidence of significant horizontal gene transfer, at least on the species level [15], *hgt* within populations may be able to explain the persistence of REPINs.

To understand how exactly *hgt* affects the evolutionary dynamics of REPIN populations, we implemented *hgt* as a simple mixing process. The *hgt* rate *h* determines the frequency at which REPINs move from one bacterium to another.

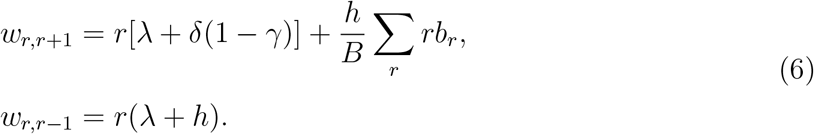

This mixing process makes the complete loss of REPINs (*b*_0_) reversible, which allows bacteria without REPINs to gain a REPIN from the rest of the population.

However, even though *hgt* provides a way to escape the zero-REPIN state, *hgt* by itself does not lead to a sustainable REPIN population. The number of REPINs in the population will still either decrease until all bacteria lose all REPINs or increase until the bacterial population is extinct.

Whether the REPIN population or the bacterial population goes extinct is mainly determined by the fatality probability *γ* for high *hgt* rates (Appendix C). For *γ* < 0.5 REPIN population size increases to infinity because REPINs successfully duplicate most of the time. In contrast, REPINs go extinct for *γ* > 0.5 due to a two-fold effect: (1) REPIN populations grow more slowly because most duplication events are unsuccessful and (2) carrying REPINs is more costly because duplication events often kill the bacterial host. Hence, *hgt* alone cannot stabilize a REPIN population in bacterial genomes

### C. Beneficial effects can lead to stable REPIN population sizes

To explain the persistence of REPINs in the genome, we propose a mutualistic relationship between REPINs and their host. In a simple model each REPIN contributes a constant benefit *α* to the host. The total fitness benefit will then be *αr*. Besides being unrealistic (adding too much of anything will eventually be detrimental), such a benefit function does not lead to a stable REPIN population. If *α* is smaller than *δ*, then the possible steady states do not change; either REPINs get purged from the genome, or the whole bacterial population goes extinct together with the REPINs. If *α* is larger than *δ*, then REPIN population size will grow to infinity as well as the bacterial population size, which is not a plausible scenario either.

Ergo the fitness benefit function needs to be more complex to describe a realistic biological scenario. An additional parameter, *w* is added to our model that modifies the beneficial effect each additional REPIN provides to the host. The following functional form changes the benefit provided by each additional REPIN, where *w* is the base of the change: [24–26],

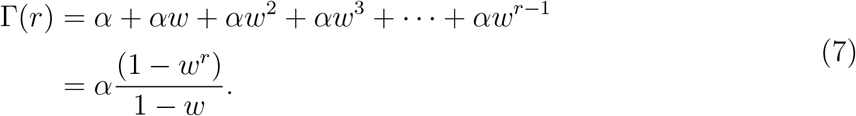

The benefit function Γ(*r*) captures the total benefit of *r* REPIN sequences (Fig. 3**A**). For *w* = 1 each REPIN provides a constant benefit *α* (discussed above). With *w* < 1, each additional REPIN provides a smaller benefit, saturating the total benefit. Similarly, with *w* > 1, each additional REPIN provides a larger benefit, exponentially increasing the total benefit. The beneficial effect of REPINs is reflected in the bacterial growth rate, *g_r_* = 1 – *B* – *rδγ* + Γ(*r*).

**FIG. 3.**
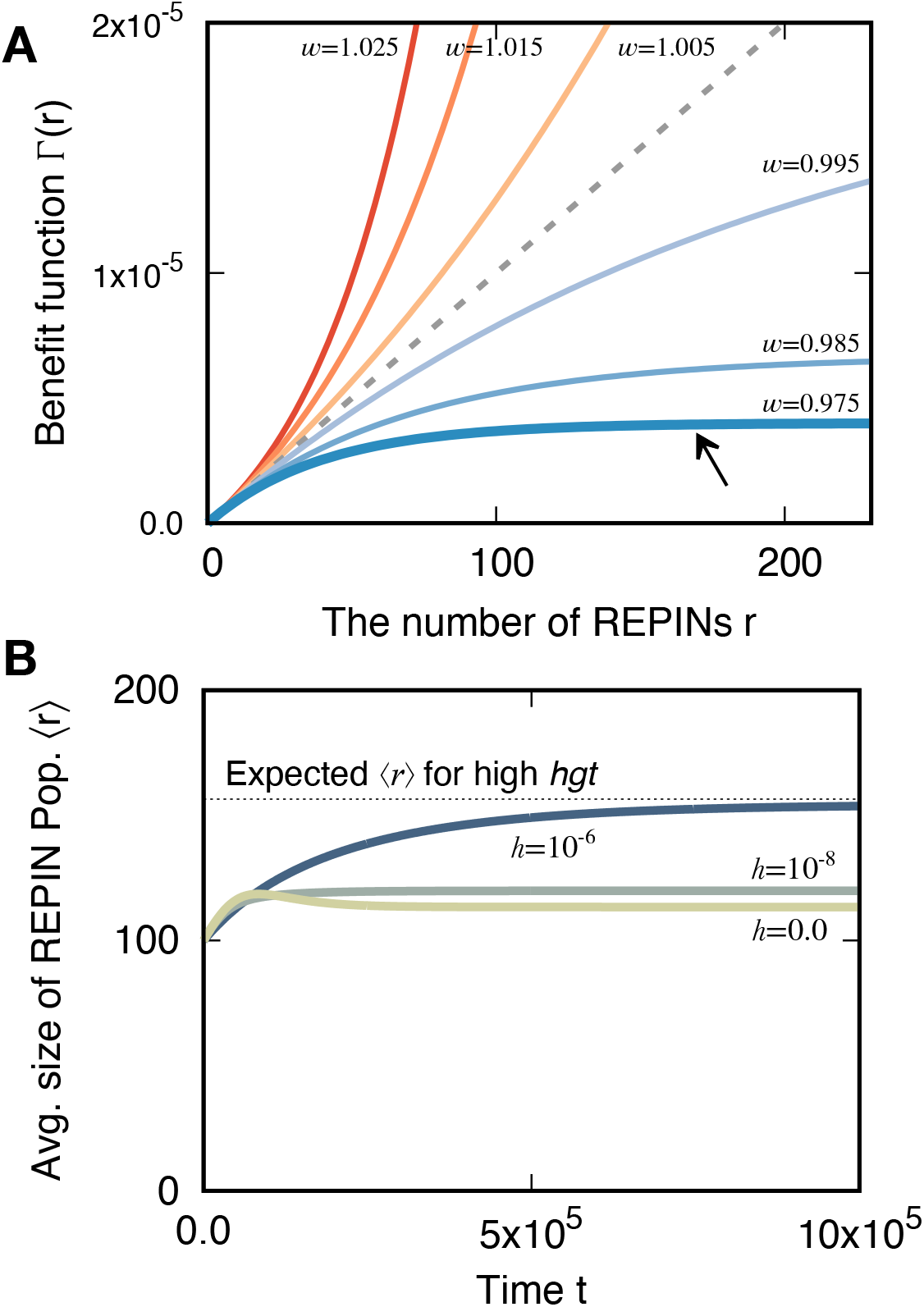
Benefit functions and dynamics of average REPIN numbers 〈*r*〉 for various *hgt* rates. **(A)** Benefit function with synergy (*w* > 1) and discounting (*w* < 1) effects. Total benefit Γ(*r*) increases with the number of REPINs *r*. With *w* = 1 the benefit a REPIN provides is constant (gray dashed line). For *w* > 1, REPIN benefits are synergistic, i.e. each additional REPIN provides a greater benefit than the previously added REPIN. For *w* < 1, REPIN benefits are discounting, i.e. each additional REPIN provides a smaller benefit than the previously added REPIN. The black arrow points at the benefit function, which is used in **(B)**. **(B)** Changes of average REPIN population sizes 〈*r*〉 over time for different *hgt* rates (h). The black dotted line is the expected REPIN population size (〈*r*〉) at the steady state for high *hgt* rates. Lower *hgt* values lead to smaller average REPIN population sizes. We used the following model parameters *γ* = 0.55, *δ* = λ = 10^−8^, *α* = 5 × 10^−8^, and *w* = 0.975.

Decreasing benefits (*w* < 1) allow a stable REPIN population to persist in the bacterial genome. For high *hgt* rates, we can analytically determine the size of the REPIN population in steady-state. To obtain a stable REPIN populations, the fatality rate needs to be high (*γ* > 0.5) and the benefit strength *α* needs to be higher than *δ*(2*γ* – 1) (Appendix D for the detailed calculation). For these conditions, we can calculate the average number of REPINs in a bacterial genome:

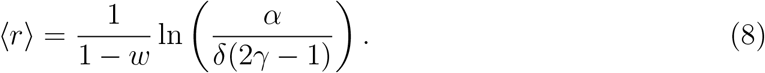

A careful analysis of the model parameters shows that there are few parameter combinations that yield a REPIN population of biologically relevant size. There are three free parameters (*α*, *γ* and *w*) that all determine the REPIN population size. We set the parameter range for *α* to 10^−7^ – 10^−9^, as this is close to the duplication rate *δ*, which also determines the fitness cost of each REPIN. The other two parameters are bounded by the model itself: *γ* can range from 0.5 < *γ* < 1 and *w* can range from 0 < *w* < 1.

For each parameter combination, we can now determine the average REPIN population size in the bacterial population. More specifically, we are interested in biologically relevant REPIN population sizes (i.e. between 91 and 323 REPINs, Tab. I). To assess, which parameter combinations yield biologically relevant REPIN populations one of the three free parameters was fixed. The other two parameters were varied across the entire range (Fig. 4).

**FIG. 4.**
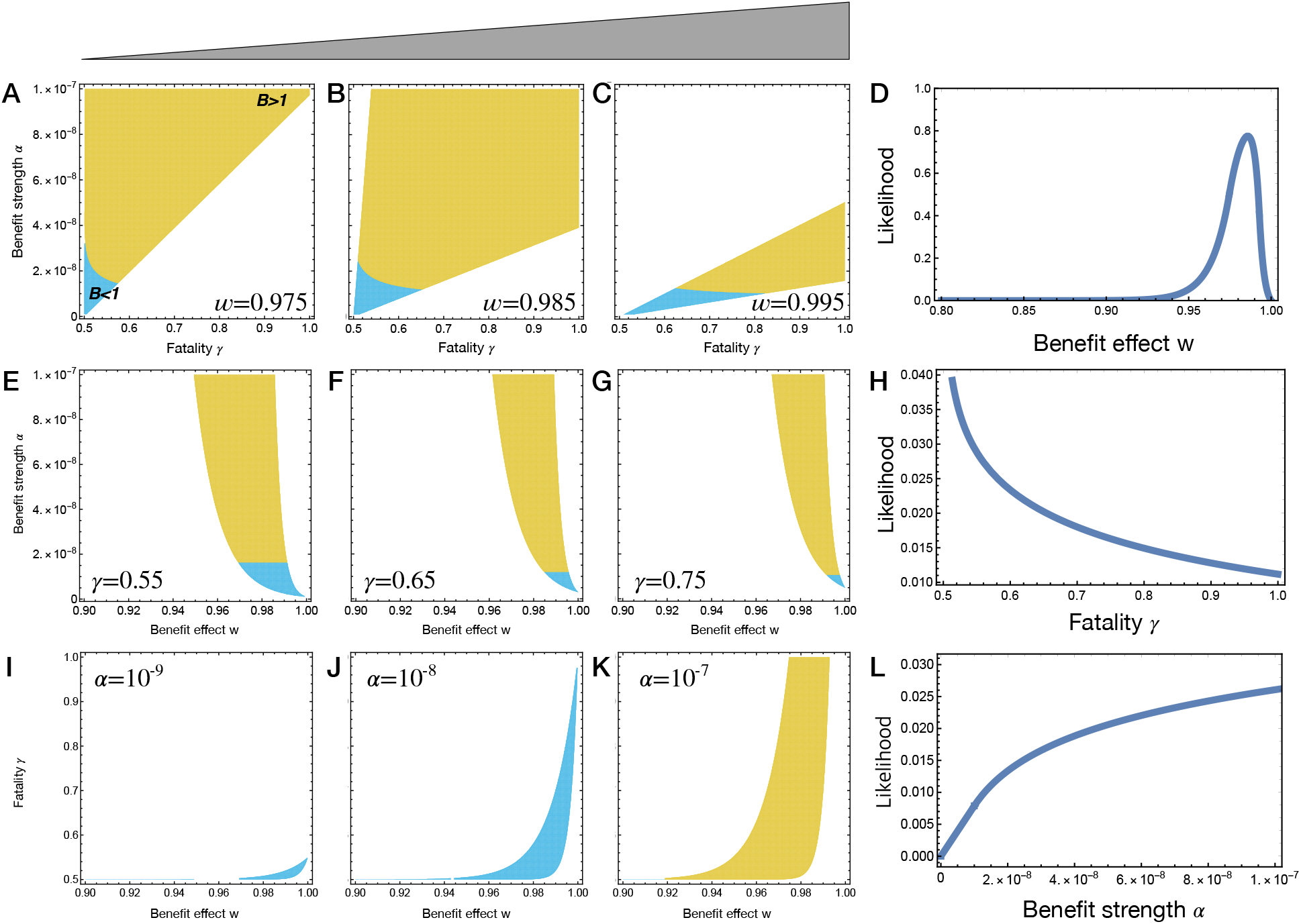
Observable parameter range and likelihood. Three model parameters, benefit effect *w*, fatality probability *γ*, and benefit strength *α*, determine the average REPIN population size 〈*r*〉. Only certain parameter combinations result in biologically relevant REPIN population sizes (i.e. between 91 and 323 REPINs, Tab. I). To visualize this observable parameter range, we fixed one parameter while the other two parameters varied. In the colored area, REPIN population sizes are between 91 and 323. The carrying capacity *K* is measured in the absence of REPINs. Hence the bacterial population size increases with REPINs in yellow colored areas, while bacterial pool size becomes smaller in blue colored areas. Each row is associated with one parameter. For example, in the first row, for three fixed benefit effects *w* we determine the observable parameter range **(A-C)**. The size of the observable parameter range is plotted in **(D)**, which corresponds to the likelihood that a *w*-value is part of a parameter combination that leads to a stable REPIN population of observable size. The second and third rows show the same plots for the fatality probability *γ* and benefit strength *α*, respectively. Note that for all parameter ranges (0 < *w* < 1, 0.5 < *γ* < 1, and 10^−9^ ≤ *α* ≤ 10^−7^) the proportion of parameter combinations that result in stable REPIN populations of observable size is only about 2%.

Without a detailed analysis of our model, the biologically relevant range of the discounting effect *w* (how strongly the benefit of each REPIN decreases with increasing REPIN number) is hard to predict. However, our model suggests that the effect needs to be in the range of 0.95 and 0.99 (Fig. 4**D**). Otherwise, the other parameter values have to become unrealistic to yield suitable average REPIN population sizes. Intuitively, this means that the benefit obtained by the host decreases by 1 – 5% with each REPIN added to the genome. Furthermore, for large discounting effects, relevant REPIN population sizes are only observed for small fatality probabilities (Fig. 4 **A**). On the contrary, small discounting effects require a small benefit strength to obtain relevant average REPIN population sizes (Fig. 4 **C**).

In our model, it is impossible to maintain a stable REPIN population if *γ* is below 0.5. Hence, at least 50% of the bacterial genome needs to be critical for long-term survival to maintain REPIN populations. Yet, a fatality probability of close to 0.5 yields the most parameter combinations to maintain a stable REPIN population (Fig. 4 **H**).

Finally, high benefit strength is most likely to yield a stable REPIN population (Fig. 4 **I-K**). Whereas low benefit strength (10^−9^) is only possible when the discounting effect is close to 1 and the fatality probability is close to 0.5 (Fig. 4 **I**).

## IV. DISCUSSION

With our study, we aim to investigate under which circumstances it is possible to maintain a self-replicating sequence population inside a bacterial genome. A simple model where REPINs duplicate in a growing bacterial population and each duplication bears a cost cannot yield a stable REPIN population. When including horizontal gene transfer to our model (mixing of REPINs between individuals of a population), the situation does not change. REPINs either die out by driving their host to extinction or are eradicated from the bacterial genome. Finally, we attached a beneficial effect to carrying REPINs. If each REPIN provides a constant beneficial effect to the bacterium, then REPIN populations are still not stably maintained. Only discounting benefits (i.e. the benefit each REPIN provides decreases with each REPIN that is added to the sequence population) can yield stable REPIN population sizes.

The specific discounting effect *w* is also quite restrictive. Only if the benefit each additional REPIN provides decreases by about 1% to 5%, are there many parameter combinations that lead to a REPIN population of relevant size (i.e. between 91 to 323 REPINs). The surprisingly narrow range of parameters will allow us to test our model in the future. In a laboratory experiment, one could, for example, delete all REPINs in a single bacterial strain (e.g. with CRISPR technology) and then add REPINs one at a time. We would expect the average additional benefit for each REPIN added to decrease by about 1 to 5%. The fitness advantage of bacteria carrying a single REPIN over bacteria carrying no REPINs should be on average in the range of the benefit strength *α*.

The benefit strength *α* is expected to be low per individual REPIN (10^−9^ < *α* < 10^−7^). Low benefit strength is a consequence of low levels of harm done by REPIN duplication due to low duplication rates (10^−8^). Interestingly, even when the benefit provided by each individual REPIN is less than the harm done (*α* < *δ*) it is possible to maintain stable REPIN populations. Furthermore, even though the average fitness of the bacteria in the population decreases the REPIN population can persist. At this point, it is unclear whether these results still hold in the absence of *hgt*. One would expect that an unrelated population without REPINs would eventually be able to outcompete resident populations. Hence, scenarios where REPINs decrease the fitness of the bacterium might ultimately not be biologically relevant since REPINs do not seem to be able to infect different bacterial species.

Another interesting aspect of our study is the result concerning the fatality probability *γ*. The fatality probability describes what proportion of REPIN duplications leads to the death of a bacterium. Our model suggests that *γ* has to be larger than 0.5 to yield a stable REPIN population. This result was somewhat surprising to us and initially did not seem to be compatible with the biological reality since studies have shown that only about 10% of all genes in the genome are essential [27, 28]. For fatality probabilities of less than 0.5 it is impossible to maintain sequence populations in bacterial genomes under our model.

One could also argue that *γ* describes the proportion of genes that are indispensable in at least one of the natural environments the bacterium encounters. That the set of genes that is essential in one environment differs from the that of a different environment has been shown in *E. coli* [29]. But of course, the experiments have been performed in the laboratory and not in the natural environment of *E. coli* and other fitness defects may also lead to eventual extinction of a bacterial population (about 49% of gene deletions have an effect on the growth phenotype). Hence, our analysis suggests that more than 50% of the bacterial genome is important enough for bacterial survival to allow stable REPIN populations to persist. If less than 50% of the bacterial genome were essential for long term survival then we would not expect to see stable REPIN populations. Instead the number of repetitive sequences should increase over time, similar to what can be observed in birds and mammals, where transposon replication is only counteracted by large genome loss events [30].

Our results in Figure 4 only hold if the horizontal gene transfer rate is much higher than the duplication rate *δ*. Active horizontal gene transfer mediated by the RAYT transposase is very unlikely to occur in nature [15]. Although REPINs and RAYTs may be passively transferred to other genomes through homologous recombination[31], the resulting rate of REPIN transfer is probably low. Hence, the results in Figure 4 might not be directly applicable to REPIN populations. Nevertheless, simulations for low horizontal gene transfer rates show that REPIN populations can persist without *hgt*, given that the REPIN population is beneficial for the host (Appendix D). In some of our simulations, we have observed stable REPIN populations that had a net negative effect on fitness. These observations were, however, rare and it is unclear whether they were a numeric artefact or possibly populations out of equilibrium.

In conclusion, our analyses show that intragenomic sequence populations cannot be stably maintained when duplications cause a fitness cost for the host. Even *hgt* is not enough to stably maintain a population of self-replicating sequences. Only when including discounting beneficial effects into our model, can it explain stable REPIN populations in bacterial genomes. Surprisingly in the presence of high *hgt* rates, the beneficial effects provided by self-replicating sequences do not have to outweigh the detrimental effects caused by REPIN duplication. Together the results of our study provide a plethora of testable hypotheses on the evolution of intragenomic sequence populations in bacterial genomes.

## APPENDICES

### Appendix A: Dynamics of the average number of REPINs

Here, we calculate the dynamics of the average number of REPINs, 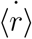,

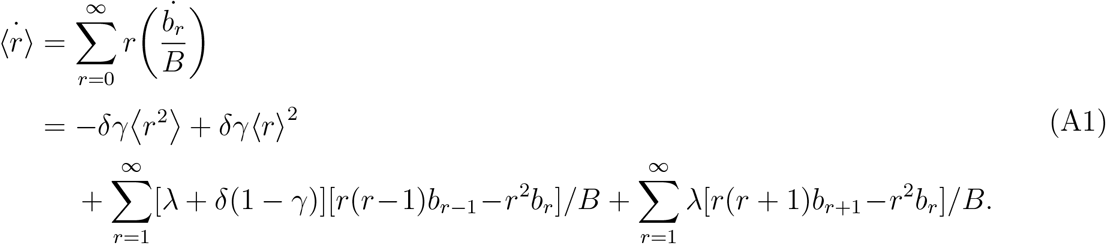

For the first summation, we obtain

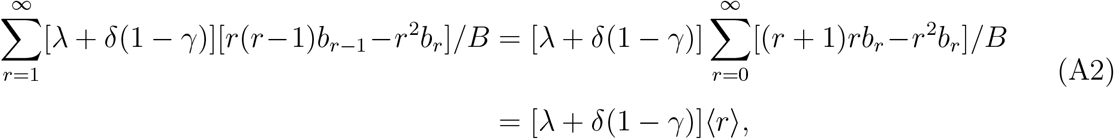

and for the second summation,

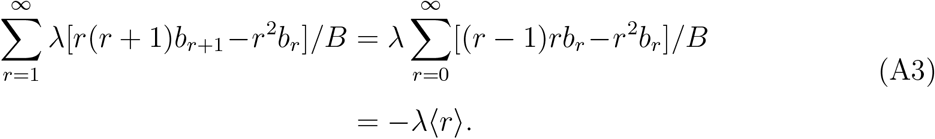

Altogether, we obtain the expression

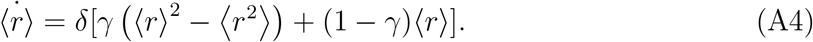

Duplication can decrease the number of REPINs by killing the host bacterium. On the other hand, successful duplication leads to an increase of REPINs. The REPIN number can increase even when this leads to lower host fitness. Precisely these two decreasing and increasing forces of REPIN numbers are reflected in the first and second terms in Eq. (A4), respectively.

### Appendix B: Trivial solutions: extinction of REPINs or extinction of both REPINs and bacteria

In this section, we will show that only trivial solutions are achieved without horizontal gene transfer and beneficial effects. For convenience, we convert the relative abundances *b_r_* into fractions *f_r_* = *b_r_/B*. Then, the main equations become,

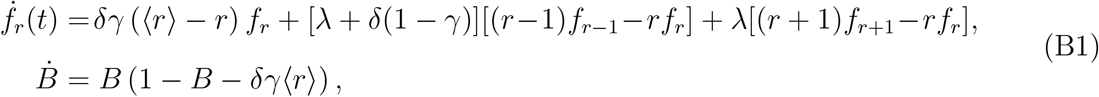

with *f_r_* = 0 for *r* < 0. From the above equation, we can obtain *f*_1_ in steady state, 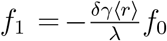. The result indicates *f*_1_ must be zero since negative values are forbidden for *f_r_*. Hence, either 〈*r*〉 = 0 or *f*_0_ = 0 must be satisfied. Then the possible solutions are the extinction of REPINs (*f*_0_ = 1 and *f_r_* = 0 for *r* > 0) or the extinction of both REPINs and bacteria (*b_r_* = 0 for all *r*).

Birth and death resulting from λ make the population diffuse in state space. If all bacteria reach the 1/(*δγ*)-REPIN state before any bacterium enters the zero-REPIN state, the whole bacterial population dies. Otherwise, REPINs will go extinct in the bacterial population, and only zero-REPIN bacteria remain.

### Appendix C: Equations of motion with *hgt*

Horizontal gene transfer can also make a bacterium lose or gain a REPIN. We assume that *hgt* happens within the population. In this case, the role of *hgt* is mixing REPINs between bacteria. No external source of REPIN is assumed to exist. With the horizontal gene transfer rate *h*, a bacterium loses REPINs proportionally to how many REPINs it contains. The insertion of REPINs can occur in any state independent of REPIN numbers. Hence, transition rates with *hgt* are given by

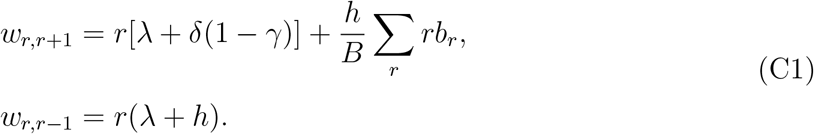

Accordingly, we obtain

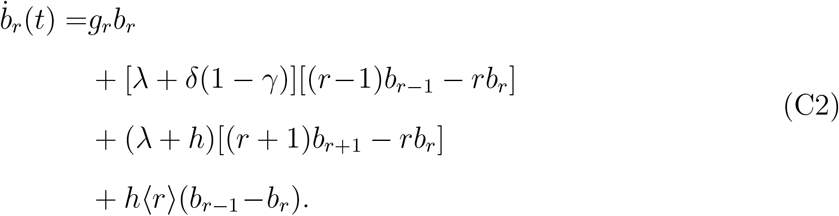

For frequencies *f_r_*, the equations become

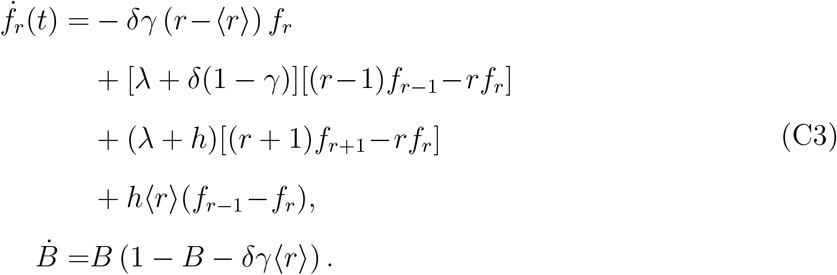

In the previous section, we found that only two trivial solutions are possible in the steady state without *hgt*. In this section, we examine possible other steady state solutions with *hgt*. Since we cannot get general solution for any *h* values, we focus on the extreme cases first and then analyse the general case.

#### a. *Low* hgt *regime*

By solving for 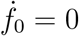 from Eq. (C3), we can obtain

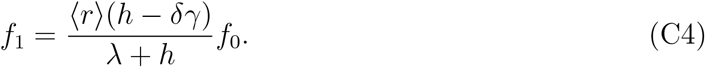

Hence, *f*_1_ should be zero for *h* ≤ *δγ* and accordingly all *f_r_* for *r* > 1 also become zero. It means that only trivial solutions are possible for *h* ≤ *δγ*.

**FIG. C1.**
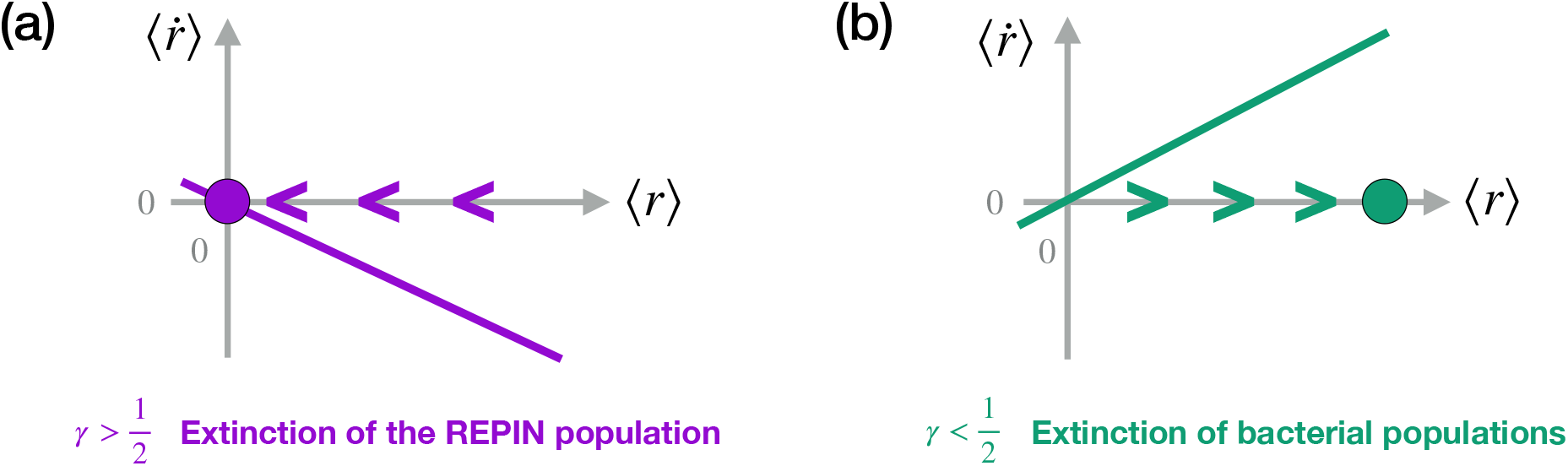
Phase portrait with high *hgt* rates. (a) For *γ* > 0.5 the number of REPINs decreases to zero. (b) For *γ* < 0.5 the number of REPINs increases without any bound, decreasing the bacterial pool *B*. Thus the whole bacterial population as well as REPINs go extinct.

#### b. *High* hgt *regime*

By solving 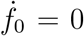 from Eq. (C3) with assumption λ, *δ* ≪ *h*, we can get

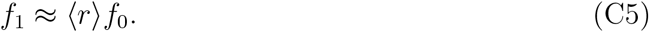

In the same way, we can recursively get the solutions,

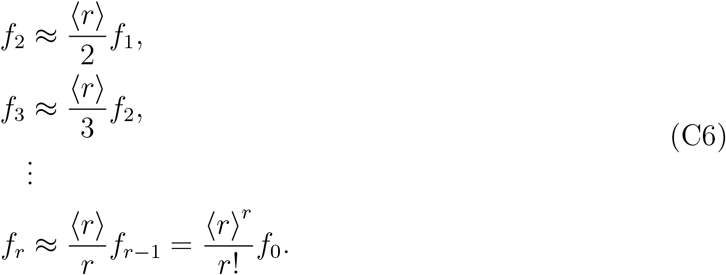

Since there is a relation between *f_r_* and 〈*r*〉, 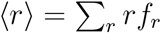, we can get *f*_0_ = *e*^−〈*r*〉^. Thus, the final solution of the stationary distribution is

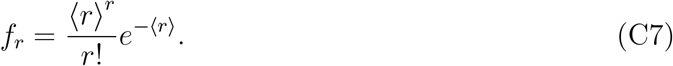

The result distribution does not depend on *h* value itself once the rate *h* is high enough.

For high *hgt* rates, the distribution is accessible so that we can calculate 〈*r*^2^〉. Plugging Eq. (C7) into Eq. (A4), we found the dynamics of the average REPIN numbers.

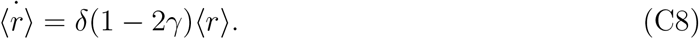

For high fatality, *γ* > 0.5, the REPIN population dies out because (1) bacteria carrying more REPINs are more likely to die than bacteria carrying fewer REPINs and (2) REPINs proliferate more slowly since most duplication are not successful (Fig. C1a). For low fatality, *γ* < 0.5, duplications are less harmful, and hence lead to increasing REPIN numbers (Fig. C1b). It happens even though an increase in REPIN numbers lead to decreasing bacterial fitness and the bacterial population eventually goes extinct. Only exactly at *γ* = 0.5 does REPIN population size 〈*r*〉 stay constant. However, this scenario is not biologically relevant because any perturbation of *γ* will lead to a population collapse.

#### c. *Intermediate* hgt *regime*

For two extreme cases, *h* ≤ *δγ* and λ ≪ *h*, we found that *hgt* cannot support the persistence of REPINs in the bacterial population. For intermediate *hgt*, *δγ* < *h* ≲ λ, we cannot apply the analytic approach, and thus we numerically investigate the possible solutions in this regime. As we shown above, the solutions of *f_r_* at the steady state can be recursively calculated from *f*_0_. For example, by solving 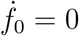, we can express *f*_1_ in terms of *f*_0_ and 〈*r*〉. In the same way, by solving 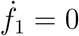, now we can express *f*_2_ in terms of *f*_1_, *f*_0_, and 〈*r*〉. Since *f*_1_ can be expressed by *f*_0_ and 〈*r*〉, *f*_2_ can be expressed by *f*_0_ and 〈*r*〉. In the same way, all *f_r_* can be expressed in terms of *f*_0_ and 〈*r*〉. Here we numerically search for a possible set of *f*_0_ and 〈*r*〉 with two constraints: (1) 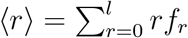 and (2) 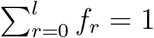. Parameter sets *γ* ∈ {0.1, 0.3, 0.5, 0.7, 0.9} and *h* ∈ {5 × 10^−9^, 10^−8^, 5 × 10^−8^, 10^−7^} with *δ* = λ = 10^−8^ and the maximum REPIN population size *l* = 50 are investigated. For all sets, only trivial solutions can be achieved, implying that *hgt* does not allow REPINs to persist.

### Appendix D: Equations of motion with mutualism

Now we explore the regime where REPINs can have a positive effect on bacterial growth,

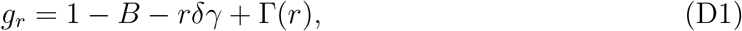

where the benefit function is

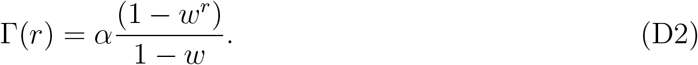

Then, the equations of motion become

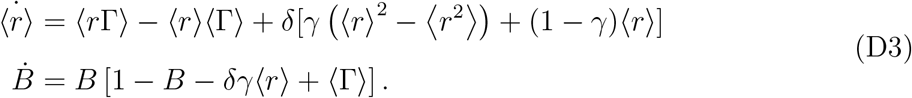

To understand the above equation, we should know the second moment of *r*, 〈*r*^2^〉. Here, we use the backward Euler method to solve Eq. (C2) with the growth rate denoted as Eq. (D1). Again we will obtain the analytic results for the high *hgt* regime first.

#### a. *High* hgt *regime*

For the high *hgt* regime, *α* ≪ *h*, mixing of REPINs between bacteria happens fast enough, and thus the distribution *f_r_* becomes smooth with a single peak around the average REPIN numbers 〈*r*〉 showing the Poisson distribution again,

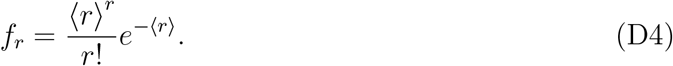

Now we can calculate 〈*r*^2^)〉 giving the solvable dynamics of 〈*r*〉,

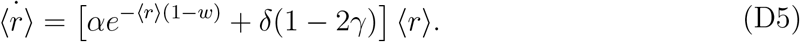

Solving the equation for an equilibrium, we can obtain the average REPIN population size 〈*r*〉 at the steady-state,

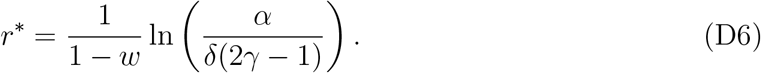

In parallel, *B* at steady-state is

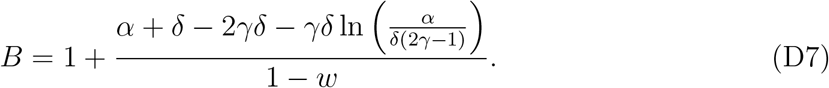

Note that this solution is valid only for *w* < 1 (discounting effect) and *γ* > 0.5. From this estimation, we can find the possible parameter ranges of *α, γ* and *w* to observe REPIN population sizes found in nature.

#### b. Stability analysis

Even if there is a fixed point for non-zero REPIN numbers, it could be unstable. In this case, a stable REPIN population cannot be maintained. Now, we will check the stability of the non-zero fixed point in Eq. (D6). The non-zero fixed point becomes stable when the 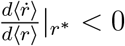. Hence, if the condition

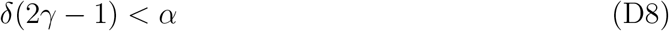

is satisfied, REPIN populations can persist at high *hgt* rates.

From Eq. (D7), the bacterial population size decreases as the result of carrying REPINs for

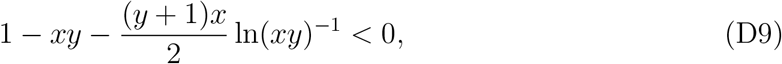

where *x* = *δ/α* and *y* = 2*γ* – 1. Surprisingly, there is a finite range in which REPINs persist even though they reduce the bacterial population size, satisfying both conditions Eqs. (D8) and (D9).

**FIG. D1.**
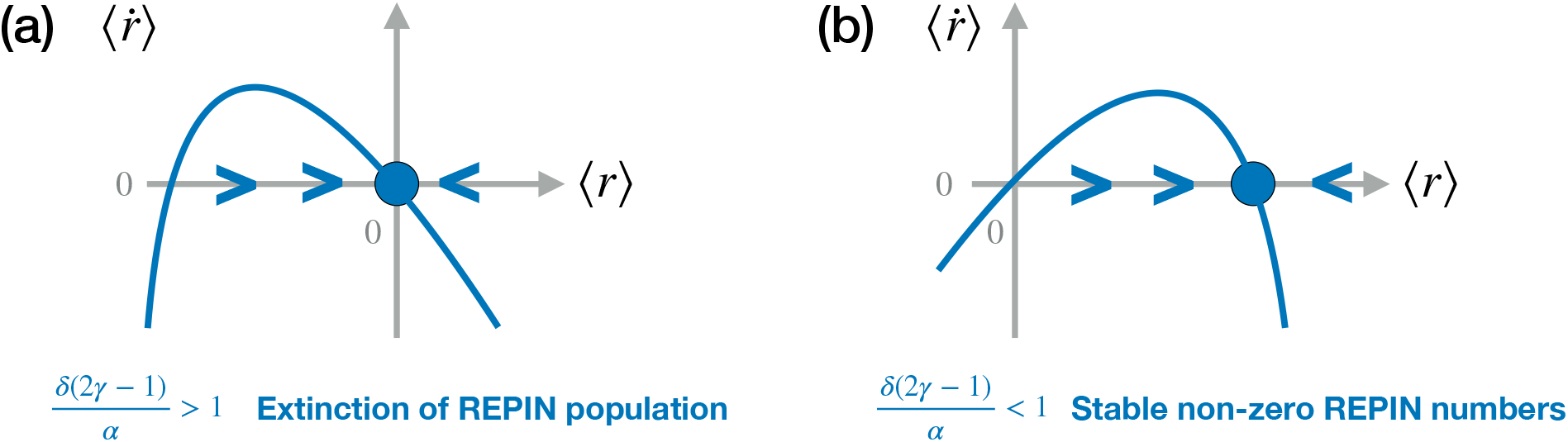
Phase portrait with high *hgt* rates and discounting beneficial effects for *γ* > 0.5. For *γ* > 0.5, duplication decreases REPIN numbers. Hence, we need a high benefit to overcome this driving force. (a) For *δ*(2*γ* – 1) > *α* the number of REPINs decreases until zero. (b) For *δ*(2*γ* – 1) < *α* a stable REPIN population can stably exist in the bacterial genome. This is even possible when the REPIN population decreases bacterial fitness.

#### c. *Low* hgt *regime*

Here, we investigate whether the expected REPIN number obtained in Eq. (D6) is a good approximation for REPIN numbers when *hgt* rates are low. We focused on the most extreme case of *h* = 0. First we randomly draw three free parameters (benefit effect *w*, fatality *γ*, and benefit strength *α*) that lead to observable REPIN numbers (91 ≤ 〈*r*〉 ≤ 323). For 500 randomly selected parameter sets, we numerically get the distribution *b_r_* at *t* = 10^6^ with an initial condition *b_r_*(0) = *δ*_*r*,100_. After obtaining numerical results, we compared the REPIN numbers obtained for *h* = 0 with values calculated for high *hgt* rates. Because REPIN numbers without *hgt* are always smaller than for high *hgt* rates, we express the difference between the REPIN numbers as a proportion (Fig. D2). 100% indicates the obtained REPIN numbers without *hgt* are the same as for high *hgt*, meaning the expression in Eq. (D6) is a good estimation. 0% means REPINs go extinct without *hgt*. When the bacterial population size exceeds the carrying capacity (carrying a REPIN population is beneficial), the REPIN population size for high *hgt* rates and no *hgt* are similar (Fig. D2, Fig. D3).

**FIG. D2.**
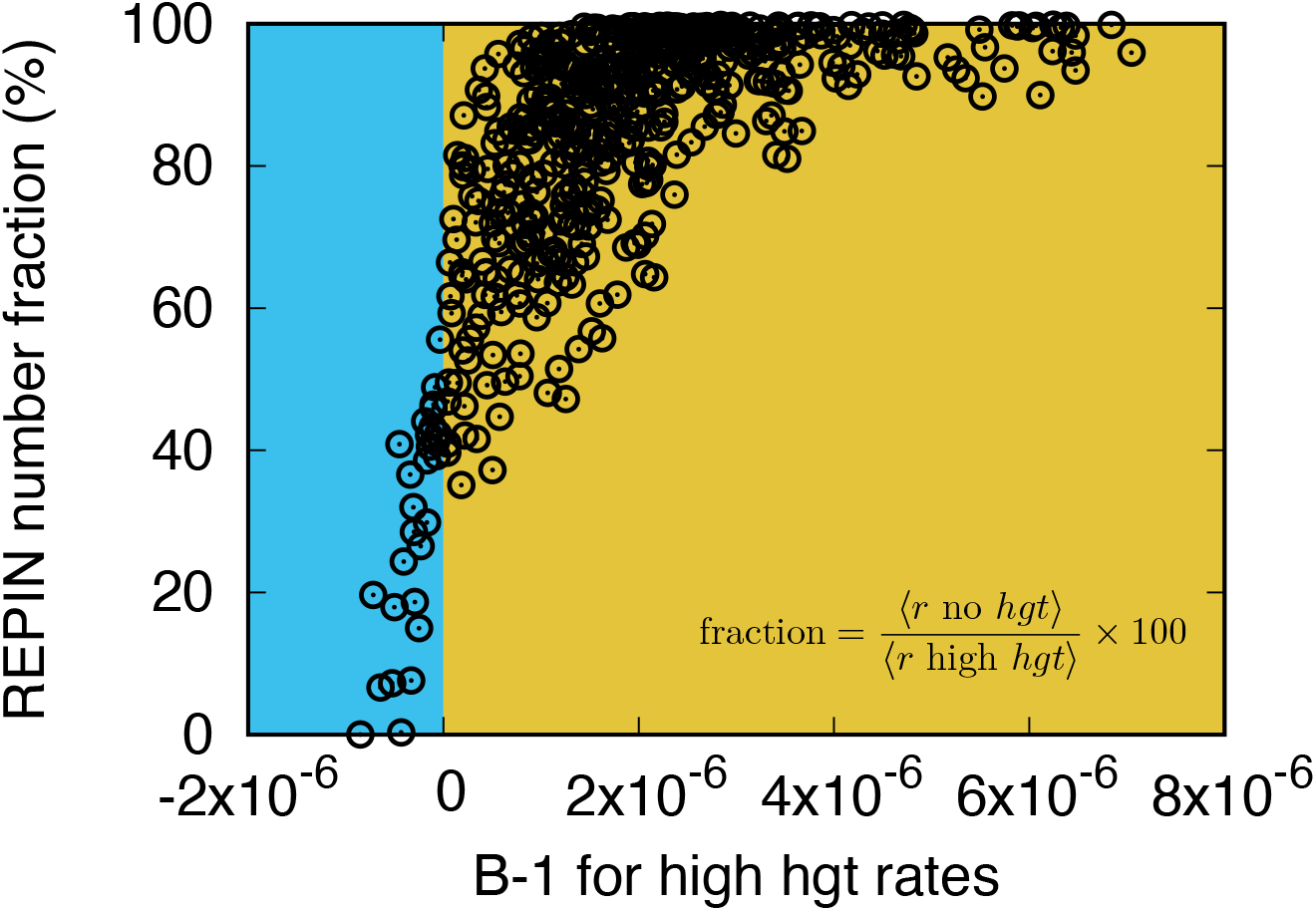
We randomly draw 500 parameter sets, which yield biologically observable REPIN numbers (91 ≤ 〈*r*〉 ≤ 323). Then, we numerically obtain REPIN numbers for *h* = 0. Y-axis shows the ratio between REPIN population sizes obtained without *hgt* and population sizes with high *hgt*. Hence, 100% indicates that the REPIN population size without *hgt* is the same as the one calculated for high *hgt*, meaning the expression in Eq. (D6) is a good estimation. The extinction of REPINs are shown as 0%. When carrying a REPIN population is beneficial for the bacterium, the results calculated for high *hgt* rates are similar to numbers obtained without *hgt* . Especially when *B* < 1, REPIN population sizes are significantly lower in the absence of *hgt* .

**FIG. D3.**
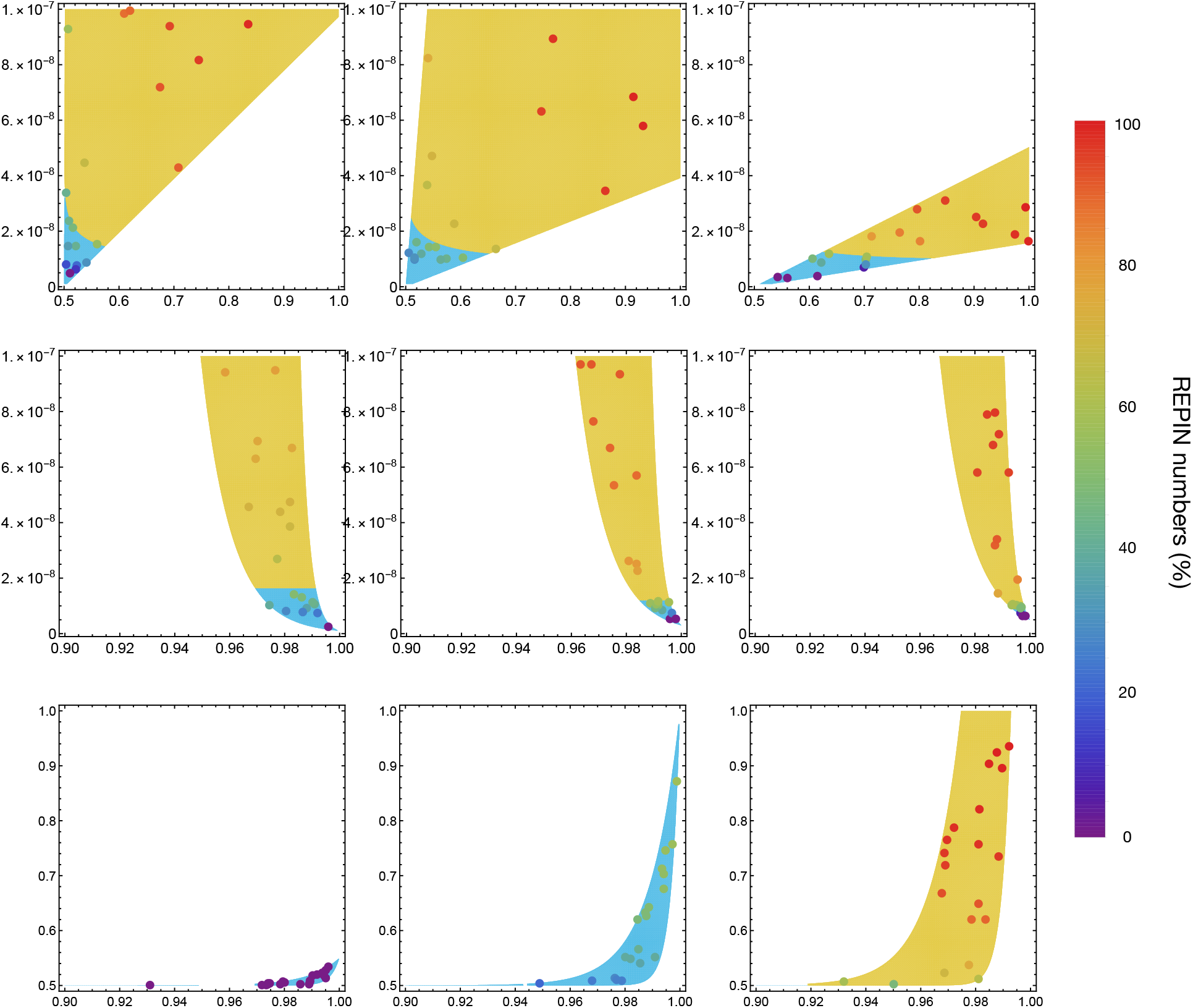
To clearly show that REPIN numbers abruptly drop when *B* < 1, we randomly sampled 20 points in each panel in Fig. 4 of the main text. If there are both regimes *B* > 1 and *B* < 1 in a single panel, we sampled 10 parameter sets for each regime. Otherwise, we sample all 20 points in one regime. We again use the same definition of REPIN number fraction in Fig. D2 to show how much REPIN numbers can be achieved compared to expected values for high *hgt* rates at given parameter sets. Points in each panel shows sampled parameter set and color indicates REPIN number fraction. As we can see, when *B* < 1 (blue colored region) is expected for high *hgt* rates, the REPIN number fractions are low. On the contrary, for *B* > 1 (yellow colored region), REPIN number fractions can reach 100%.

